# Network analysis of workshop activities reveals increasing transdisciplinarity of German biodiversity research community

**DOI:** 10.1101/2020.02.26.966432

**Authors:** Jonas Geschke, Martina Lutz, Katrin Vohland

**Author notes:** This research has been conducted within and about the activities of the Network-Forum on Biodiversity Research Germany (NeFo). NeFo is implemented by the Helmholtz-Centre for Environmental Research GmbH – UFZ in Leipzig and the Museum für Naturkunde - Leibniz Institute for Evolution and Biodiversity Science in Berlin. From February 2009 to August 2019, NeFo was funded by the Federal Ministry of Education and Research (BMBF). Many thanks to Marianne Darbi, Nike Sommerwerk, Rainer Schliep and Graham Prescott for providing very constructive comments on the manuscript. Supplementary material: The dataset and all files needed to re-run the R analysis presented in this paper will be made available through doi.org/10.5281/zenodo.3550542 once the final, peer-reviewed paper is published.

## Abstract

Boundary spanning activities in the biodiversity science-policy interface are urgently needed. Effective science communication and uptake of scientific findings by policymakers is crucial for a successful, cross-scale policy implementation. For this, national platforms promoting knowledge exchange between different stakeholder groups are key. Established in 2009, the Network-Forum on Biodiversity Research Germany (NeFo) until 2018 has organized more than 40 workshops bringing together actors from science, policy and society. In this paper, we present a network and cluster analysis of these NeFo workshops. Based on this, we discuss the importance of science-policy interface projects and networks as knowledge brokers and boundary organizations, as well as challenges in using network analysis as a tool for evaluating workshop impacts. Based on the network analysis outcomes as well as experiences in the conduction of workshops, recommendations to strengthen the innovation impact of networking efforts are drawn.

## 1. Introduction

Decision makers and actors at the science-policy interface increasingly acknowledge the importance of biodiversity and the risks arising from its loss (e.g. FAO, 2019; IPBES, 2019). The growing number of both national and international policy strategy documents during the past decade as well as the growing number of grant programs and the establishment of institutions active at the science-policy interface illustrate this change towards more biodiversity awareness and inter- and transdisciplinary biodiversity research. Nevertheless, the link between scientific knowledge and political action still has to be strengthened and a number of proposals have been made for improving communication and networking between different actors (e.g. Shanley and López, 2009; Turnhout et al., 2016). In order to adequately support different biodiversity-related policy processes, more integrative research institutions and efforts towards inter- and transdisciplinary research are needed (Mehring et al., 2017).

The Network-Forum on Biodiversity Research Germany (NeFo) was initially founded as a project in 2009 and has ever since been established as a brand at the biodiversity science-policy interface (with three project phases, running from February 2009 to December 2012, January 2013 to July 2014 and August 2014 to August 2019). Its main goal was to bring together relevant actors from biodiversity research in Germany, in order to strengthen the inter- and transdisciplinarity of the German biodiversity research institutions and scientists and at the same time foster their dialogue with politics, administration and practice (Marquard et al., 2011). In order to do so, NeFo has intensively worked on establishing contacts and a network spanning different disciplines of biodiversity research. Over the past ten years, NeFo brought together experts from science, policy and non-governmental organizations, which led to controversial, intensive and productive discussions. During its development process, NeFo had different thematic focusses: While it supported the development of biodiversity research as an independent interdisciplinary research field first, it also addressed different thematic core areas at the science-policy interface. Thereby, NeFo became a key player in structuring and communicating the processes and outcomes of the Intergovernmental Science-Policy Platform on Biodiversity and Ecosystem Services (IPBES) in Germany and fostering a high participation of German scientists in IPBES assessments. Next to its networking activities with respect to different biodiversity research topics (e.g. land use, monitoring, ecosystem services, pollination, synthetic biology, etc.), NeFo initiated different capacity building formats in support of IPBES, which are now annually organized by the German IPBES Coordination Office (the National IPBES Forum in Germany) or co-organized by multiple national science-policy platforms throughout Europe (the Pan-European IPBES Stakeholder Consultations (PESC); FRB, 2017).

Given the diverse and fragmented landscape of biodiversity research institutions recorded in the “NeFo Research Atlas” (Chamsai et al., 2011; Vohland et al., 2012; Schliep et al., 2016) and the potential incompleteness of the atlas hindering a robust social network analysis (Geschke and Vohland, 2018), we sought to analyse the role of workshops and expert talks (hereafter “workshops”) for inter- and transdisciplinarity within the German biodiversity research community and beyond. Therefore, in this paper, we discuss the impact of the workshops and expert talks (hereafter “workshops”) organized by NeFo in the years from 2010 to 2018 as a tool for networking, science communication and knowledge exchange tool. We demonstrate how NeFo has developed as an interface between science, policy and society. Throughout the paper, we first introduce the methodology of social network analysis as well as the dataset and statistical parameters used. Subsequently, we present and interpret the main results from the network analysis and discuss limitations of the data and methodology and conclude how further work can improve the outcome, efficiency and strategic development of inter- and transdisciplinary networking efforts.

## 2. Methodology and data

### 2.1 Social network analysis

Social networks are comprised of different actors and the links between them (Wasserman and Faust, 1994). Graphically, social networks are represented by nodes (for the actors) and lines (for the links). The links can be analysed based on directionality (directed, e.g. for the flow of resources or information, or undirected) and on weight (e.g. for the value of resources or the number of collaborative publications) (Wasserman and Faust, 1994). The overall pattern of a social network can be described and visualized more detailed by including statistical values such as the density of a network (percentage of total possible links realized), the degree centrality (number of links an individual actor has) or betweenness centrality (measure for how often an individual actor is part of the shortest path between two other actors that are disconnected themselves) (Wasserman and Faust, 1994; Hawe et al., 2004; Fuhse, 2016). With such parameters, which are among the mostly used parameters for the evaluation of social networks (Lang and Leifeld, 2008), different relationships can be visualized (see e.g. Brockhaus et al., 2014; Hauck et al., 2015). Betweenness centrality, for example, often is used to evaluate the power of actors (Granovetter, 1983; Krackhardt, 1990; Melbeck, 2004). Next to classic “one-mode” networks with nodes of the same type (e.g. people or institutions), “two-mode” networks describe networks with nodes of two different types (e.g. people and events) (Hawe et al., 2004; Luke, 2015; Fuhse, 2016).

Social network analyses have proven to be a useful tool to assess and visualize the effects of social activities (e.g. Lang and Leifeld, 2008; Varone et al., 2017; Giurca and Metz, 2018). The effects of knowledge exchange tools such as web portals and workshops are hardly measurable, especially in policy advice and innovation support. Here, social network analyses provide a variety of tools and statistics that can be used to create indicators for the quality and development of networks (Posner and Cvitanovic, 2019). Also, existing networks can be assessed in order to derive strategic starting points towards the future developments of a network and of human-nature-relations (e.g. Bodin and Crona, 2009; Newig et al., 2010; Calvet-Mir et al., 2015; Salpeteur et al., 2017).

### 2.2 Dataset

All files needed for replicating this social network analysis are provided in the supplementary material (R.zip). The dataset matrix, which was put together from the participant lists of workshops organized by NeFo, contained anonymous data for how many representatives of an institution attended a certain workshop, with each row containing the institution and country where its representative(s) came from and each column representing a different workshop. Thus, in the visualizations, the nodes represent institutions and workshops and the lines represent the institution’s participation in a workshop. For better understanding of the patterns and developments within the biodiversity research community, institutions were divided into the categories science, public authority, business, NGO, media, education, church and “others/ no category”. The latter category consisted of people who did not declare any institution.

The final dataset consisted of a total of 42 workshops organized from 2010 to 2018 (table 1) with a total of 1204 participants representing 485 institutions from 70 countries (figure 1); 79 participants did not declare an institution.

**Table 1:**
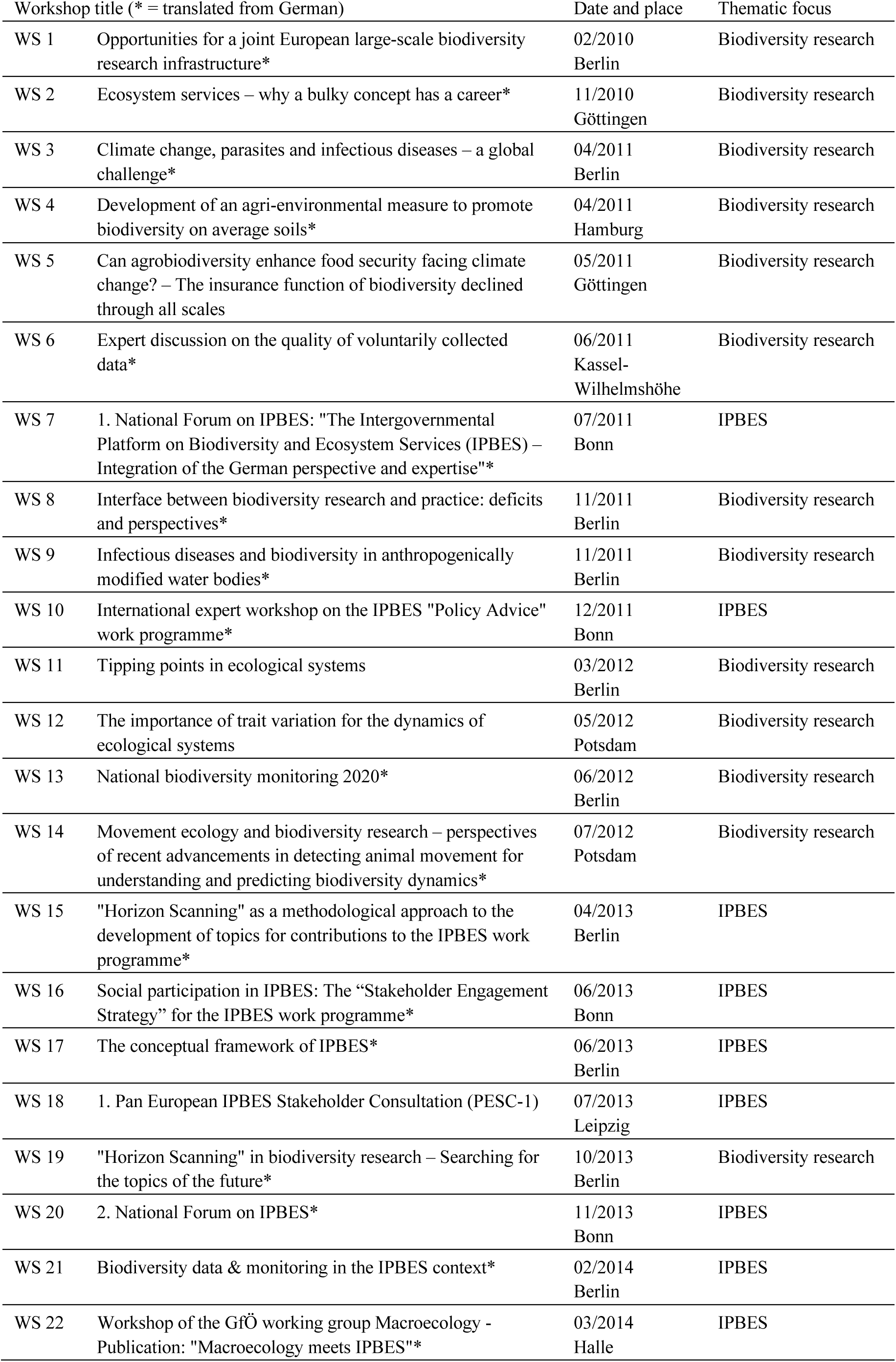
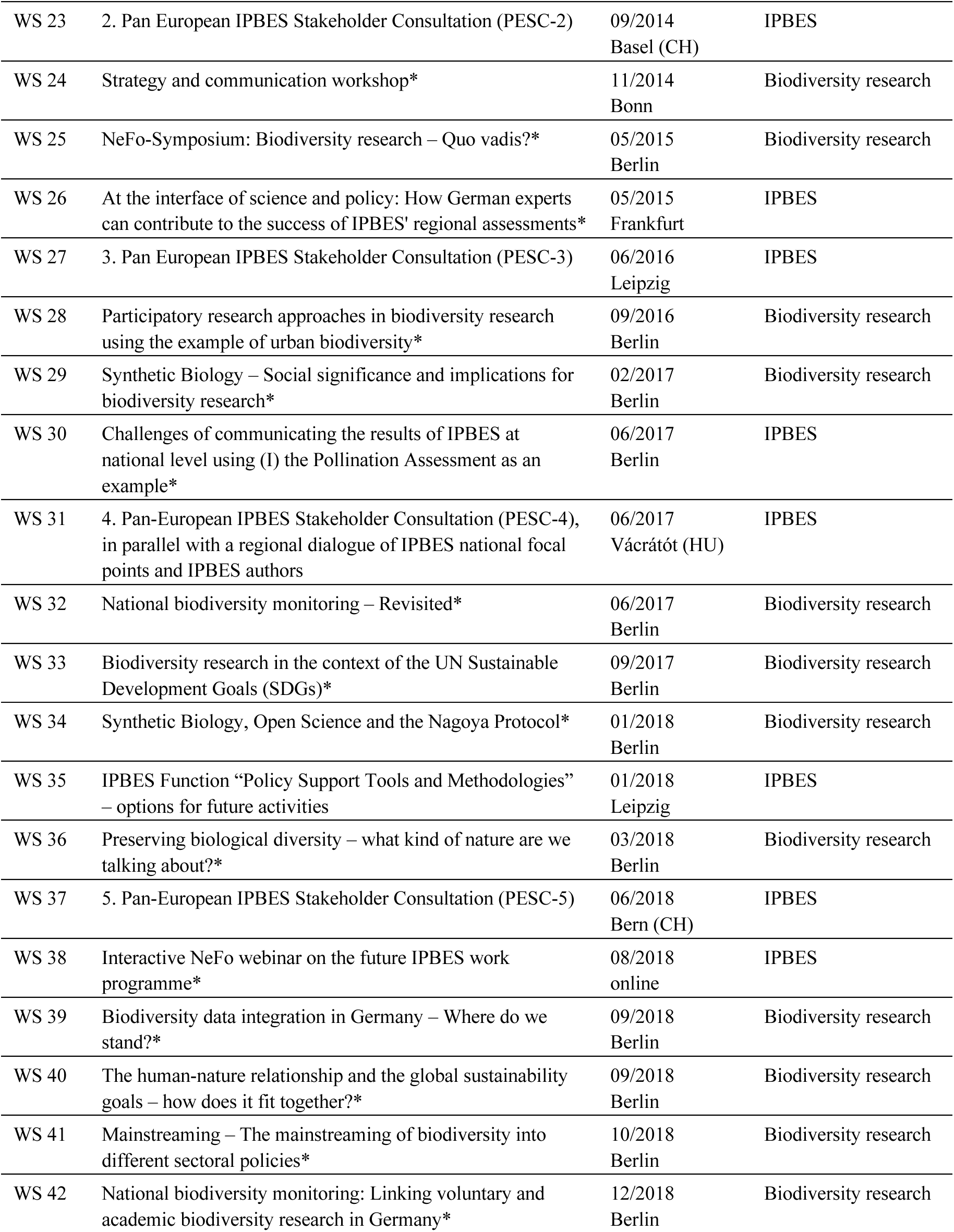
Overview of the workshops organized by NeFo from 2010 to 2018.

| Workshop title (* = translated from German) |  | Date and place | Thematic focus |
| --- | --- | --- | --- |
| WS 1 | Opportunities for a joint European large-scale biodiversity research infrastructure* | 02/2010<br>Berlin | Biodiversity research |
| WS 2 | Ecosystem services – why a bulky concept has a career* | 11/2010<br>Göttingen | Biodiversity research |
| WS 3 | Climate change, parasites and infectious diseases – a global challenge* | 04/2011<br>Berlin | Biodiversity research |
| WS 4 | Development of an agri-environmental measure to promote biodiversity on average soils* | 04/2011<br>Hamburg | Biodiversity research |
| WS 5 | Can agrobiodiversity enhance food security facing climate change? – The insurance function of biodiversity declined through all scales | 05/2011<br>Göttingen | Biodiversity research |
| WS 6 | Expert discussion on the quality of voluntarily collected data* | 06/2011<br>Kassel-Wilhelmshöhe | Biodiversity research |
| WS 7 | 1. National Forum on IPBES: "The Intergovernmental Platform on Biodiversity and Ecosystem Services (IPBES) – Integration of the German perspective and expertise"* | 07/2011<br>Bonn | IPBES |
| WS 8 | Interface between biodiversity research and practice: deficits and perspectives* | 11/2011<br>Berlin | Biodiversity research |
| WS 9 | Infectious diseases and biodiversity in anthropogenically modified water bodies* | 11/2011<br>Berlin | Biodiversity research |
| WS 10 | International expert workshop on the IPBES "Policy Advice" work programme* | 12/2011<br>Bonn | IPBES |
| WS 11 | Tipping points in ecological systems | 03/2012<br>Berlin | Biodiversity research |
| WS 12 | The importance of trait variation for the dynamics of ecological systems | 05/2012<br>Potsdam | Biodiversity research |
| WS 13 | National biodiversity monitoring 2020* | 06/2012<br>Berlin | Biodiversity research |
| WS 14 | Movement ecology and biodiversity research – perspectives of recent advancements in detecting animal movement for understanding and predicting biodiversity dynamics* | 07/2012<br>Potsdam | Biodiversity research |
| WS 15 | "Horizon Scanning" as a methodological approach to the development of topics for contributions to the IPBES work programme* | 04/2013<br>Berlin | IPBES |
| WS 16 | Social participation in IPBES: The "Stakeholder Engagement Strategy" for the IPBES work programme* | 06/2013<br>Bonn | IPBES |
| WS 17 | The conceptual framework of IPBES* | 06/2013<br>Berlin | IPBES |
| WS 18 | 1. Pan European IPBES Stakeholder Consultation (PESC-1) | 07/2013<br>Leipzig | IPBES |
| WS 19 | "Horizon Scanning" in biodiversity research – Searching for the topics of the future* | 10/2013<br>Berlin | Biodiversity research |
| WS 20 | 2. National Forum on IPBES* | 11/2013<br>Bonn | IPBES |
| WS 21 | Biodiversity data & monitoring in the IPBES context* | 02/2014<br>Berlin | IPBES |
| WS 22 | Workshop of the GfÖ working group Macroecology - Publication: "Macroecology meets IPBES"* | 03/2014<br>Halle | IPBES |
| WS 23 | 2. Pan European IPBES Stakeholder Consultation (PESC-2) | 09/2014<br>Basel (CH) | IPBES |
| WS 24 | Strategy and communication workshop* | 11/2014<br>Bonn | Biodiversity research |
| WS 25 | NeFo-Symposium: Biodiversity research – Quo vadis?* | 05/2015<br>Berlin | Biodiversity research |
| WS 26 | At the interface of science and policy: How German experts can contribute to the success of IPBES' regional assessments* | 05/2015<br>Frankfurt | IPBES |
| WS 27 | 3. Pan European IPBES Stakeholder Consultation (PESC-3) | 06/2016<br>Leipzig | IPBES |
| WS 28 | Participatory research approaches in biodiversity research using the example of urban biodiversity* | 09/2016<br>Berlin | Biodiversity research |
| WS 29 | Synthetic Biology – Social significance and implications for biodiversity research* | 02/2017<br>Berlin | Biodiversity research |
| WS 30 | Challenges of communicating the results of IPBES at national level using (I) the Pollination Assessment as an example* | 06/2017<br>Berlin | IPBES |
| WS 31 | 4. Pan-European IPBES Stakeholder Consultation (PESC-4), in parallel with a regional dialogue of IPBES national focal points and IPBES authors | 06/2017<br>Vácrátót (HU) | IPBES |
| WS 32 | National biodiversity monitoring – Revisited* | 06/2017<br>Berlin | Biodiversity research |
| WS 33 | Biodiversity research in the context of the UN Sustainable Development Goals (SDGs)* | 09/2017<br>Berlin | Biodiversity research |
| WS 34 | Synthetic Biology, Open Science and the Nagoya Protocol* | 01/2018<br>Berlin | Biodiversity research |
| WS 35 | IPBES Function “Policy Support Tools and Methodologies” – options for future activities | 01/2018<br>Leipzig | IPBES |
| WS 36 | Preserving biological diversity – what kind of nature are we talking about?* | 03/2018<br>Berlin | Biodiversity research |
| WS 37 | 5. Pan-European IPBES Stakeholder Consultation (PESC-5) | 06/2018<br>Bern (CH) | IPBES |
| WS 38 | Interactive NeFo webinar on the future IPBES work programme* | 08/2018<br>online | IPBES |
| WS 39 | Biodiversity data integration in Germany – Where do we stand?* | 09/2018<br>Berlin | Biodiversity research |
| WS 40 | The human-nature relationship and the global sustainability goals – how does it fit together?* | 09/2018<br>Berlin | Biodiversity research |
| WS 41 | Mainstreaming – The mainstreaming of biodiversity into different sectoral policies* | 10/2018<br>Berlin | Biodiversity research |
| WS 42 | National biodiversity monitoring: Linking voluntary and academic biodiversity research in Germany* | 12/2018<br>Berlin | Biodiversity research |

**Figure 1.**
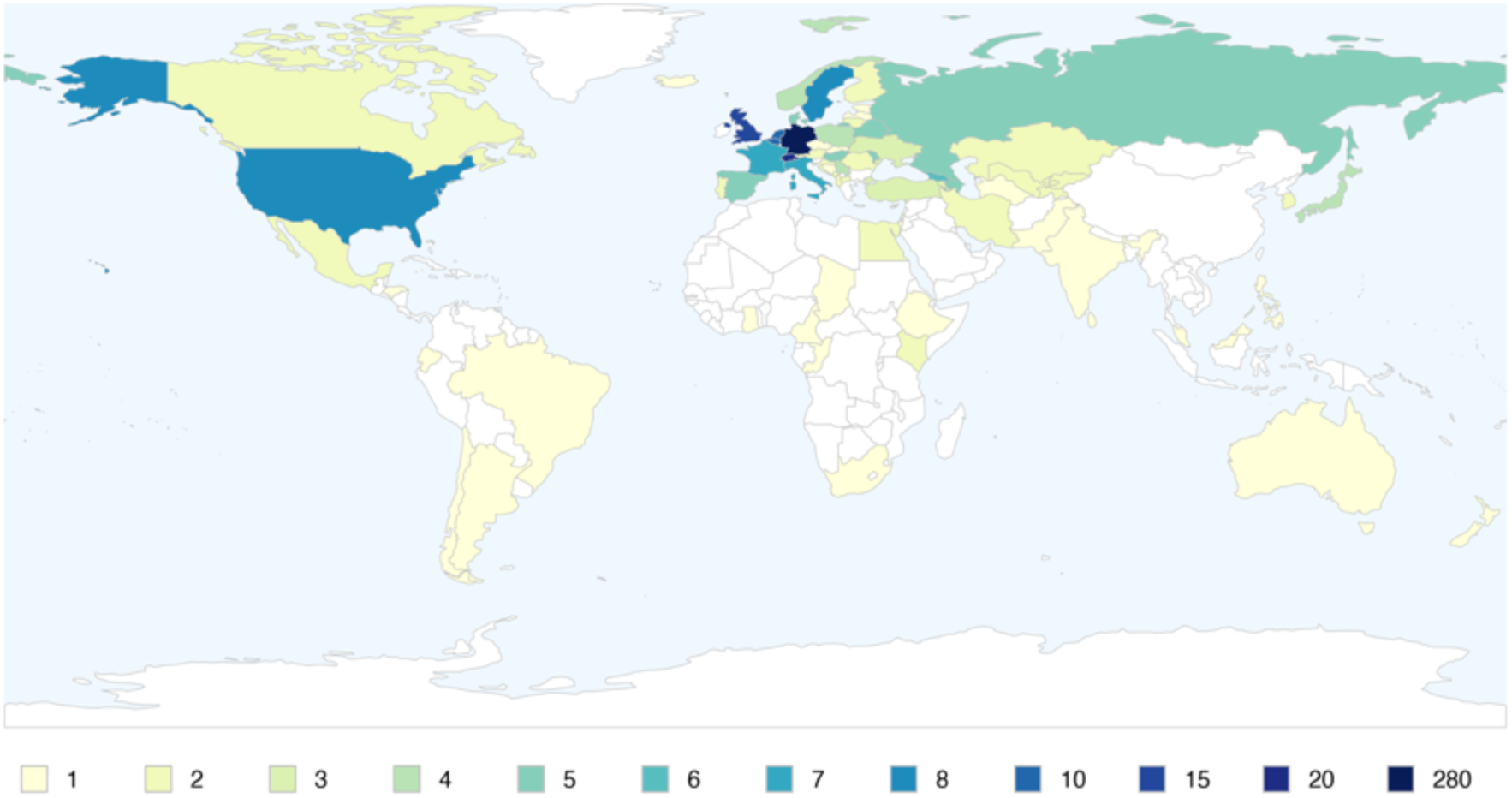
Global map of where the representatives from institutions participating in NeFo workshops from 2010 to 2018 came from. The colours represent the number of institutions from a certain country. The dataset was comprised of 485 institutions from 70 countries.

### 2.3 Statistical parameters and cluster analysis

The results section consists of two parts: the presentation of the degree of participation of the different institutional categories in the workshops (section 3.1 Participation trends over time), including a general linear model (GLM) analysis of the participation of the institutional categories over time and a network analysis (section 3.2 Cluster analysis). For clarity, network visualizations are provided as “two-mode” network. The betweenness centrality, however, is processed as classic “one-mode” network, in order to remove bias from workshop nodes in between the institution nodes (i.e. the workshops were taken as the links between institutions and not as extra nodes between the institutions). The degree and betweenness values of the institutions are provided in the supplementary material (network_statistics.csv). All links were calculated without direction and weight (meaning that an institution node always represents at least one participant from the institution). Instead, the total number of participants per institution is illustrated by colour. The cluster analysis was run based on a modified approach of the propagating labels algorithm developed by Raghavan et al. (2007). For details on this, check the R script in the supplementary material.

## 3. Results and their interpretation

Based on the NeFo workshops from 2010 to 2018, this case study drafts the complexity of the biodiversity research community in Germany. Both the network and cluster analysis of the workshops clearly illustrate their role for inter- and transdisciplinary networking as well as science communication and knowledge exchange at the science-policy-society interface. This is described in more detail:

### 3.1 Participation trends over time

As shown in table 2, the institutional categories “media”, “education”, “church” and “others/ no category” make up a small proportion of the relative attendance data only. We therefore focused the analysis on the following categories: “science”, “public authority”, “business” and “NGOs”. All 42 workshops had science institutions participating; 39 workshops had NGO institutions, 36 had public authority institutions and 26 had business institutions participating. On mean average, 66 % of the participating institutions came from science. The *absolute* number of scientific institutions remained stable over the years. A GLM calculation of the *relative* participation of the different categories (figure 2; table 3) however revealed a significant decrease in percentage of science institutions over time (p = 0.03 *), while the percentage of institutions from business was significantly increasing (p = 0.03 *). A GLM of the absolute numbers of participating institutions per categories (see table 3) resulted in a significant increase of institutions from NGOs over time (p = 0.01 *).

**Table 2:** Percentages of institutions participating in NeFo workshops per category (rounded).

| Workshop | Institutional categories |  |  |  |  |  |  |  |
| --- | --- | --- | --- | --- | --- | --- | --- | --- |
|  | Science | Public authority | Business | NGOs | Media | Education | Church | Others/ no category |
| WS 1 | 85 | 7 | 2 | 7 | - | - | - | - |
| WS 2 | 86 | 2 | 9 | - | 2 | - | - | - |
| WS 3 | 78 | 12 | 5 | 5 | - | - | - | - |
| WS 4 | 50 | 35 | 10 | 5 | - | - | - | - |
| WS 5 | 94 | - | - | 6 | - | - | - | - |
| WS 6 | 58 | 8 | - | 33 | - | - | - | - |
| WS 7 | 45 | 20 | - | 10 | - | - | - | 25 |
| WS 8 | 56 | 6 | - | 39 | - | - | - | - |
| WS 9 | 88 | 6 | - | 3 | - | - | - | 3 |
| WS 10 | 32 | 53 | - | 13 | 1 | - | - | 1 |
| WS 11 | 89 | 4 | - | 7 | - | - | - | - |
| WS 12 | 100 | - | - | - | - | - | - | - |
| WS 13 | 66 | 20 | 1 | 11 | 1 | - | - | - |
| WS 14 | 100 | - | - | - | - | - | - | - |
| WS 15 | 78 | 11 | 6 | 6 | - | - | - | - |
| WS 16 | 54 | 15 | 8 | 23 | - | - | - | - |
| WS 17 | 87 | 7 | - | 7 | - | - | - | - |
| WS 18 | 58 | 21 | 1 | 19 | - | - | - | 1 |
| WS 19 | 70 | - | 17 | 13 | - | - | - | - |
| WS 20 | 67 | 17 | 11 | 5 | - | - | - | - |
| WS 21 | 80 | 11 | - | 9 | - | - | - | - |
| WS 22 | 92 | 4 | - | 2 | - | - | - | 2 |
| WS 23 | 38 | 31 | 8 | 23 | - | - | - | - |
| WS 24 | 53 | 16 | 11 | 21 | - | - | - | - |
| WS 25 | 56 | 15 | 7 | 15 | 1 | - | - | 4 |
| WS 26 | 70 | 17 | 7 | 7 | - | - | - | - |
| WS 27 | 65 | 15 | - | 20 | - | - | - | - |
| WS 28 | 78 | 5 | 5 | 8 | - | - | - | 3 |
| WS 29 | 47 | 22 | 6 | 14 | 6 | - | - | 6 |
| WS 30 | 60 | 15 | 5 | 15 | 5 | - | - | - |
| WS 31 | 43 | 38 | 3 | 16 | - | - | - | - |
| WS 32 | 81 | 8 | 6 | 6 | - | - | - | - |
| WS 33 | 69 | 6 | 6 | 19 | - | - | - | - |
| WS 34 | 61 | 19 | 3 | 16 | - | - | - | - |
| WS 35 | 74 | 22 | - | 4 | - | - | - | - |
| WS 36 | 25 | 12 | 31 | 13 | 2 | - | - | 16 |
| WS 37 | 58 | 27 | - | 15 | - | - | - | - |
| WS 38 | 45 | 10 | 15 | 20 | - | - | - | 10 |
| WS 39 | 76 | 6 | 15 | 3 | - | - | - | - |
| WS 40 | 21 | 4 | 10 | 14 | - | 6 | 1 | 45 |
| WS 41 | 69 | - | 9 | 22 | - | - | - | - |
| WS 42 | 63 | - | - | 37 | - | - | - | - |

**Table 3:** Results of the GLM calculations of participating institutions per category.

| Institutional category | Relative number of institutions per category |  |  | Absolute number of institutions per category |  |  |
| --- | --- | --- | --- | --- | --- | --- |
|  | Estimate | Std. Error | p | Estimate | Std. Error | p |
| Science | -0.54 | 0.24 | 0.03 * | -0.02 | 0.16 | 0.90 |
| Public authority | -0.01 | 0.15 | 0.92 | 0.00 | 0.11 | 0.95 |
| Business | 0.18 | 0.08 | 0.03 * | 0.15 | 0.08 | 0.07 |
| NGOs | 0.20 | 0.12 | 0.10 | 0.12 | 0.06 | 0.01 * |
| Media | 0.01 | 0.02 | 0.54 | 0.01 | 0.01 | 0.53 |
| Education | 0.02 | 0.01 | 0.13 | 0.02 | 0.01 | 0.13 |
| Church | 0.00 | 0.00 | 0.13 | 0.00 | 0.00 | 0.13 |
| Others/ no category | 0.14 | 0.10 | 0.17 | 0.19 | 0.10 | 0.06 |

**Figure 2.**
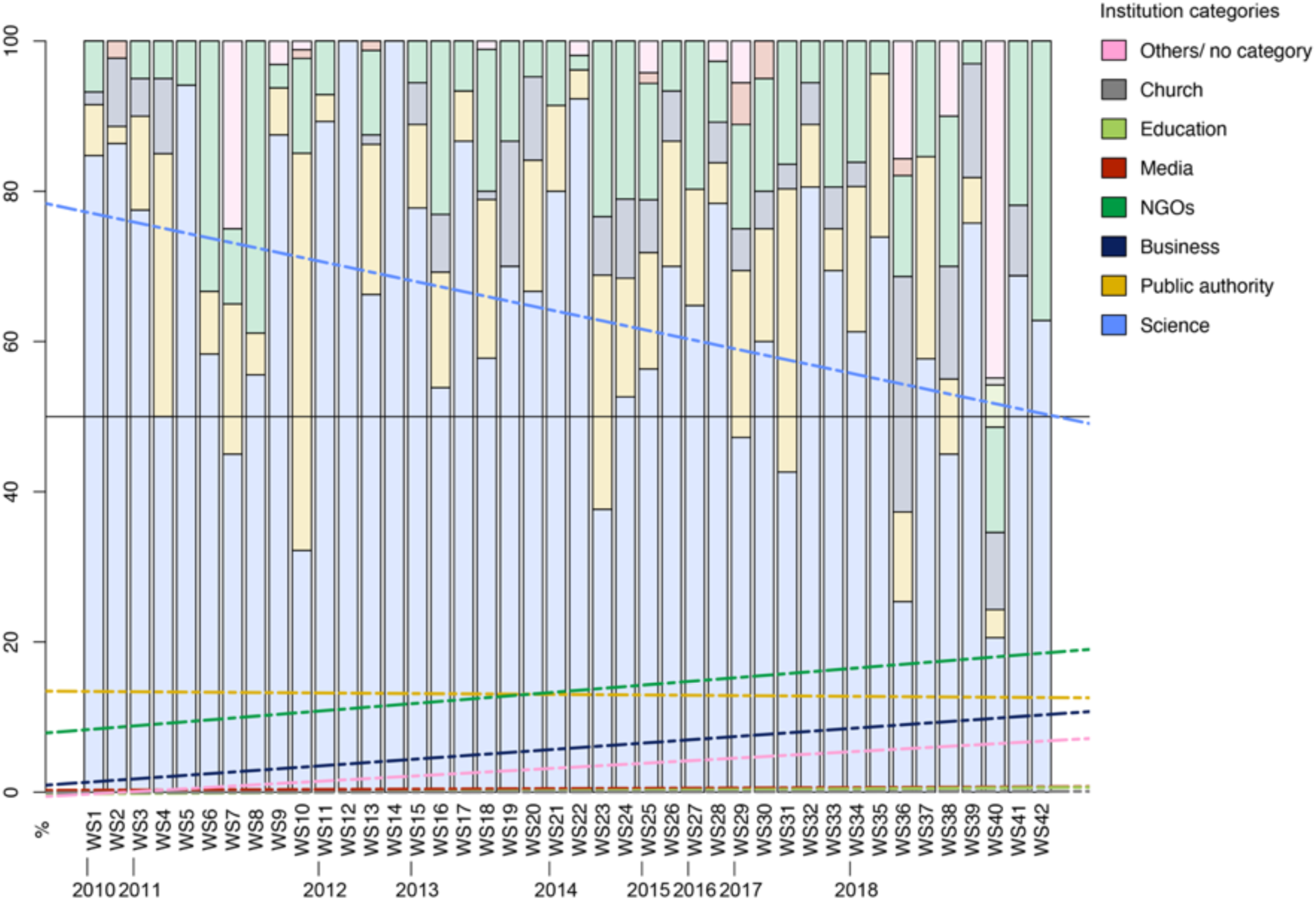
Relative participation and GLM trends of institutional categories in NeFo workshops from 2010 to 2018.

This allows the interpretation that the workshops extended the NeFo network across disciplines, in particular with business and NGO institutions, while science institutions (the key target group of NeFo) remained the key members the network.

### 3.2 Cluster analysis

The network of workshops and participating institutions (figure 3) illustrates that the central area of the network is dominated by science institutions as well as a number of institutions from national public authorities and NGOs. The peripheral areas are dominated by international public authorities on the one hand and business as well as by the “others/ no category” category with no institutions declared on the other hand. This observation is validated by a cluster analysis that resulted in nine communities (figure 4) that can be grouped into three core clusters that correspond to the three project phases of NeFo:

1. Activating the biodiversity research community: The network’s central community represents the German biodiversity science-policy interface community. This community is the key target group of NeFo and was particularly addressed through workshops that have been conducted in the first project phase.
2. Supporting IPBES: As part of the second project phase, the project’s focus was on the development of the IPBES process and the engagement of IPBES stakeholders in IPBES work. Workshop 10 and workshops 18, 23, 27, 31 and 37 represent the policy targeted NeFo activities through IPBES-related workshops (e.g. FRB, 2017). Note that the workshop 10 community is separated from the others, as workshop 10 was held to support the conceptualization of IPBES prior to its establishment while workshops 18, 23, 27, 31 and 37 contributed to the stakeholder engagement after the establishment of IPBES.
3. Opening up biodiversity research towards societal engagement: Lastly, workshops 36 and 40 represent NeFo activities in the format of evening events that were conducted to reach out to actors from civil society and get them engaged in biodiversity and sustainability related discussions. The conduction of evening events as such did not begin until 2018, as part of the third project phase.

**Figure 3.**
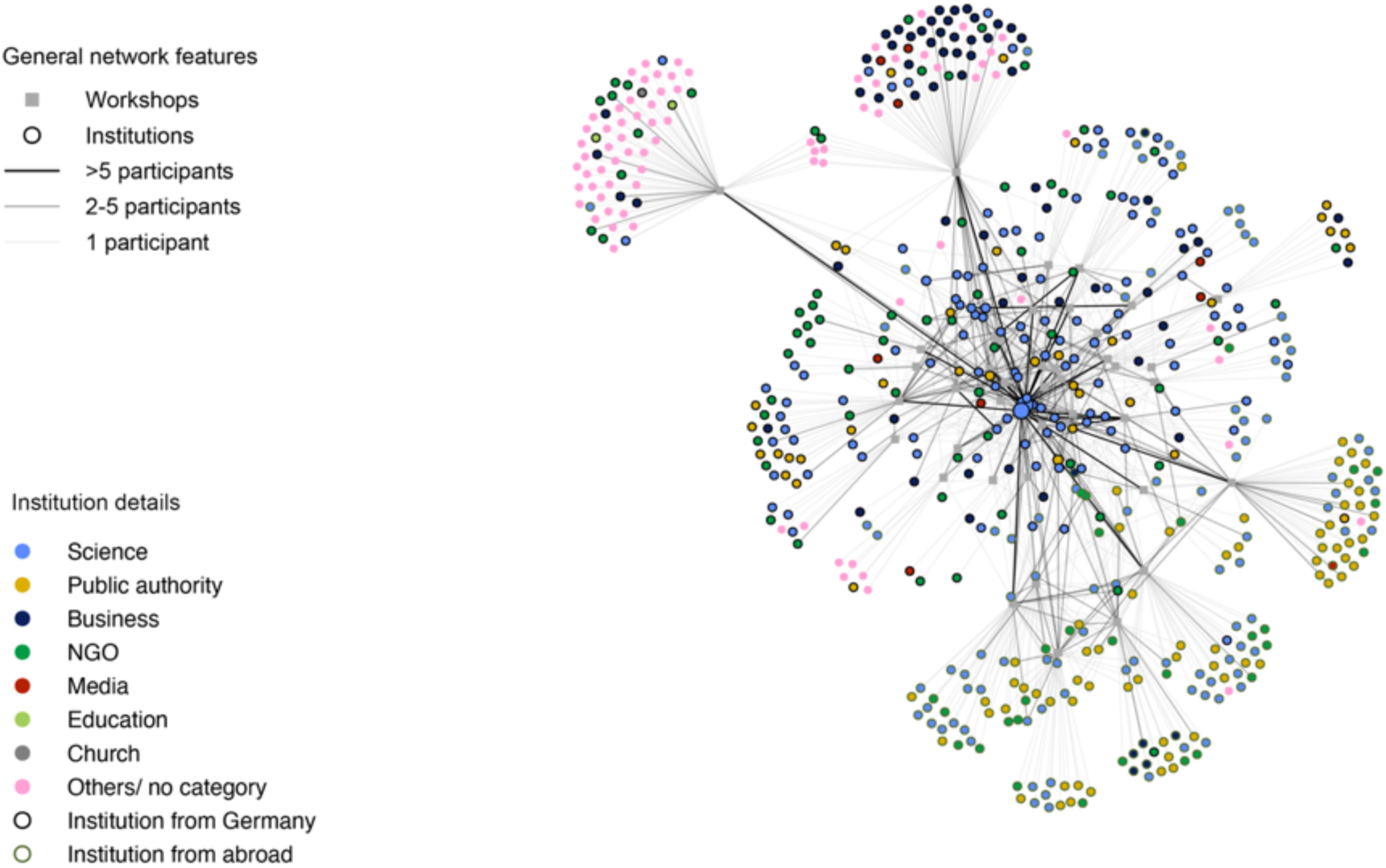
Network graph of the 42 NeFo workshops from 2010 to 2018. The institution nodes are coloured according to their assigned category, with their size reflecting the betweenness centrality of an institution; the institution node borders represent the country of origin of the institutions; the lines are coloured according to the number of participants from the institutions in a workshop, the participant number is not considered within any line statistics.

**Figure 4.**
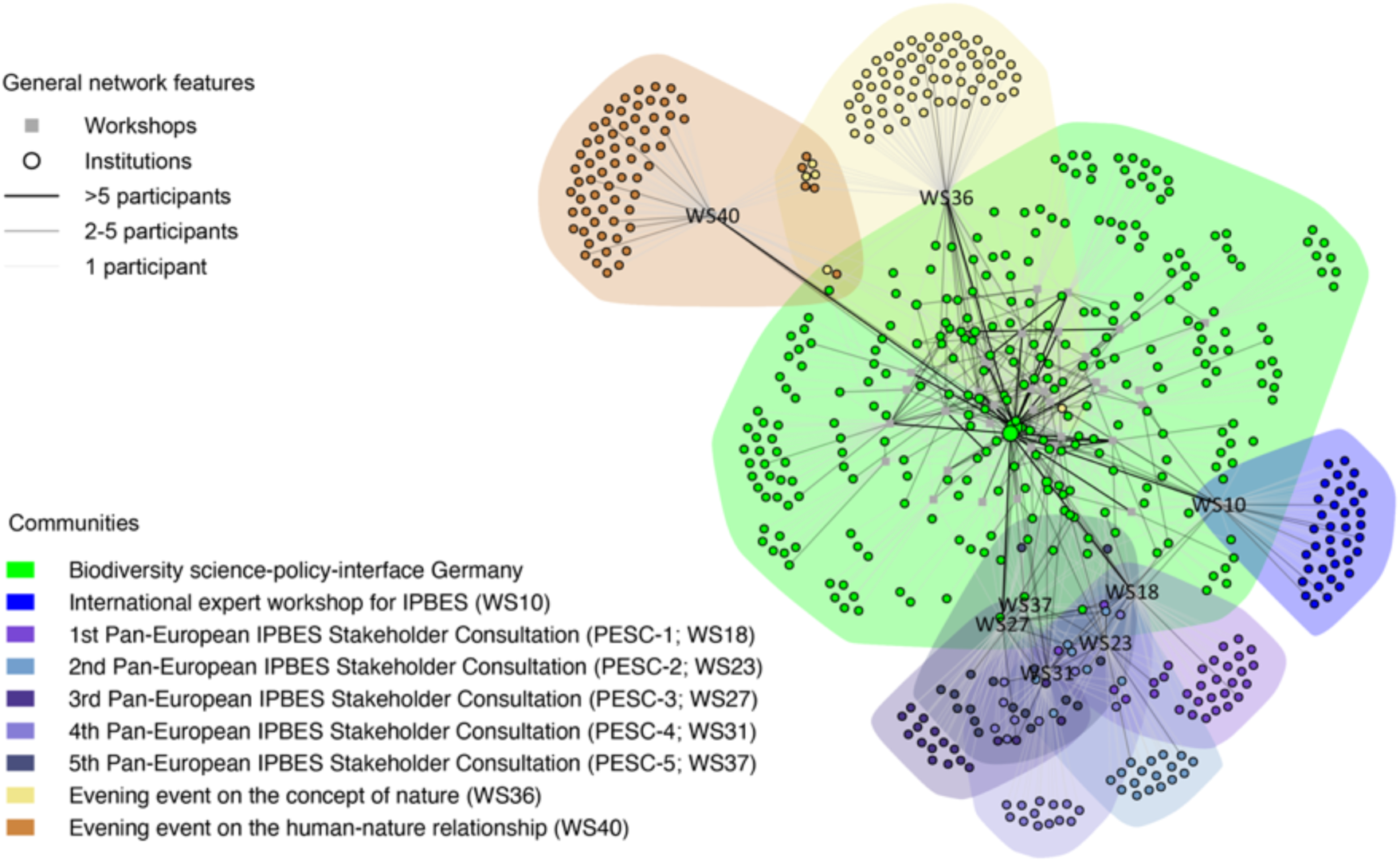
Cluster graph of the 42 NeFo workshops from 2010 to 2018. The institution nodes are coloured according to their modelled community, with their size reflecting the betweenness centrality of an institution; the lines are coloured according to the number of participants from the institutions in a workshop, the participant number is not considered within any line statistics.

Therefore, when considering the time axis of the workshops, respectively the three project phases of NeFo, the cluster analysis shows that NeFo has developed from (1) a national science-policy interface to (2) an international IPBES-focused science-policy interface and finally to (3) a science-policy-society interface. For the work as science-policy-society interface, different workshop and engagement formats are shown to be essential for bringing science, policy and society closer to biodiversity related topics, foster discussions and raise awareness.

## 4. Discussion of the data and methodology

Social network analyses often have certain methodological limitations that are mainly caused by the network boundaries given by the data used, as one can have different approaches to gather network data and every change in nodes or links affects the network analysis outcome (Fuhse, 2016). In this analysis, the data represents institutions attending workshops – without a statistical weight for how many participants represented an institution and without an assessment of who actually interacted with whom. This network analysis therefore does not illustrate the personal networks of participants but linkages between institutions. In future studies, an analysis of project collaborations could potentially complement the dataset and analysis presented. However, as the workshop participants actively came to the workshops upon invitation by NeFo, the analysis allows an assessment of the community developments throughout the NeFo project phases. Here, the results show an increasing integrativity of NeFo workshops during the past decade: The absolute number of science institutions participating did not change significantly, while there was a significant increase of participation of business and NGO institutions – which led to a reduction in relative numbers of science institutions participating in NeFo workshops. This trend of the biodiversity research getting more integrative has been hypothesized earlier (Reuter et al., 2015; Schliep et al., 2016) and now is demonstrated by this quantitative network analysis. Important to note is that the interpretation of network analysis results needs qualitative knowledge about the network features and habits. Also, a network analysis and especially cluster analysis can be run with a variety of algorithms, which may bring different results. Therefore, the methodological approach presented in this paper is just one possible perspective on the NeFo workshops and their transdisciplinary networking impact.

As outlined earlier, due to their small degree of participation, the categories media, education, church and others were not considered further in the evaluation of the results. The category “others/ no category” cannot be further specified, as the participants did not declare any institution. However, the increasing number of people intentionally not declaring (and thus representing) any institution suggests that there is an increasing interest in the private and societal sector in participating in biodiversity research discourses.

## 5. Conclusions

### 5.1 NeFo as knowledge broker and boundary organization

The overall picture of NeFo workshops shows an inter- and transdisciplinary science-policy-society interface with a focus on biodiversity relevant discourses. What does this tell about the function of a project such as NeFo for the biodiversity research community in Germany? Complex knowledge systems such as the biodiversity research community in Germany need actors who provide links between institutions and policy processes. Only with effective, transparent and credible knowledge brokers at the interfaces between different disciplines and societal sectors, decision-makers can be informed and different perspectives comprehensively integrated (Neßhöver et al., 2016; Morin et al., 2017; Sarkki et al., 2019). Network analyses can help to assess knowledge exchange flows within a given network and identify or evaluate knowledge brokers (Toikka, 2010; Crona and Parker, 2011; Weiss et al., 2012; Cvitanovic et al., 2017), as demonstrated by the presented analysis.

More generally, NeFo can be considered a boundary organization, providing the opportunity to connect different boundaries or social worlds, such as science, policy and society (Star and Griesemer, 1989). Boundary organizations stand at the intersection of one or more boundaries and enable participation from all sides of the boundaries they deal with (Guston, 1999, 2001). By actively addressing boundary objects or topics and offering space for interdisciplinary and intersectoral discussions, NeFo aimed to overcome certain boundaries that hinder effective knowledge exchange. In this process, knowledge brokers play the important role of facilitators (Bednarek et al., 2018). Knowledge brokers can achieve different boundary spanning impacts, such as improved knowledge exchange, more diverse and stronger social networks, increased trust, empowered scientists, the creation of policy windows to link knowledge production with use in policy making and enhanced capacity of policy makers and their institutions (Posner and Cvitanovic, 2019). As demonstrated by our analysis, NeFo was able to improve the inter- and transdisciplinary knowledge exchange in the German biodiversity research community and, over time, increased the diversity of sectors involved. With its workshops and studies (e.g. FRB, 2017; Schliep et al., 2018), NeFo also assisted in and evaluated the establishment of a national coordinating system for German scientists that wanted to get involved in IPBES – therefore empowered scientists and provided capacity building activities for both scientists and decision-makers. In consideration of the performance measures presented by Gustafsson and Lidskog (2018) and Posner and Cvitanovic (2019), which are adaptiveness, competence and the achievement of the above mentioned boundary spanning impacts, we argue that NeFo is successfully acting as a boundary organization in the German biodiversity research and policy sector.

### 5.2 General conclusions and workshop recommendations

The methodology of social network analysis is a promising approach for visualizing, assessing and strategically evaluating networking activities. However, network analyses of complex scientific networking efforts require analyses at multiple scales, including not only the institutional level but also the levels of projects and personal networks. Such levels need to be equally assessed in order to be able to evaluate the whole picture of networking efforts. This is especially crucial due to the fact that institutionalized networks potentially are based upon personal networks. Therefore, the data base for a social network analysis needs to be as comprehensive and clearly defined as possible and should consider multiple dimensions, e.g. network scale, space, time and robust network boundaries. To complement this case study, further analyses considering project-based collaboration between biodiversity research institutions are needed. Also, further analyses should be backed up by qualitative information, e.g. gathered through surveys and interviews. Nevertheless, the on-hand case study reveals that knowledge brokers and boundary organizations such as NeFo can integrate actors who have been isolated so far and widen the inter- and transdisciplinary knowledge exchange. It illustrates that national science-policy interface projects have the potential to play an important role in bringing together different actors from different disciplines and attract actors from outside the core scientific sector. As such, from this social network analysis, we draw the following recommendations to strengthen the innovation impact of workshops as networking efforts and thus boundary spanning activities:

- Workshops should address not only well-established topics as boundary objects but also topics that are known but highly complex and not fully understood cross-sectoral.
- Workshops should be organized in cooperation with other actors (by creating a collaborative boundary organization to conduct a workshop), in order to reach out to a wider, potentially new audience and increase the diversity in disciplines and sectors.
- Actors that have a broker role within a social network analysis should be particularly asked for future cooperation, as they have the potential to integrate so far largely isolated actors and sub-networks.
- Actors that are new to the network should be kept in discourse, e.g. through targeted invitation or integration into follow-up networking efforts.
- Depending on the workshop format, appropriate engagement and follow-up opportunities should be given to all participants.

